# Intramolecular interactions dominate the autoregulation of *Escherichia coli* stringent factor RelA

**DOI:** 10.1101/680231

**Authors:** Kathryn Jane Turnbull, Ievgen Dzhygyr, Søren Lindemose, Vasili Hauryliuk, Mohammad Roghanian

## Abstract

Amino acid starvation in *Escherichia coli* activates the enzymatic activity of the stringent factor RelA, leading to accumulation of the alarmone nucleotide (p)ppGpp. The alarmone acts as an intercellular messenger to regulate transcription, translation and metabolism to mediate bacterial stress adaptation. The enzymatic activity of RelA is subject to multi-layered allosteric control executed both by ligands – such as ‘starved’ ribosomal complexes, deacylated tRNA and pppGpp – and by individual RelA domains. The auto-regulation of RelA is proposed to act either *in cis* (inhibition of the enzymatic activity of the N-terminal region, NTD, by regulatory C-terminal region, CTD) or *in trans* (CTD-mediated dimerization leading to enzyme inhibition). In this report, we probed the regulatory roles of the individual domains of *E. coli* RelA and our results are not indicative of RelA dimerization being the key regulatory mechanism. First, at growth-permitting levels, ectopic expression of RelA CTD does not interfere with activation of native RelA, indicating lack of regulation *via* inhibitory complex formation in the cell. Second, in our biochemical assays, increasing RelA concentration does not decrease the enzyme activity, as would be expected in the case of efficient auto-inhibition *via* dimerization. Third, while high-level CTD expression efficiently inhibits the growth, the effect is independent of native RelA and is mediated by direct inhibition of protein synthesis, likely *via* direct interaction with the ribosomal A-site. Finally, deletion of the RRM domain of the CTD region leads to growth inhibition mediated by accumulation of (p)ppGpp, suggesting de-regulation of the synthetic activity in this mutant.

## 1 Introduction

Bacteria have numerous mechanisms for sensing and adapting to stressful conditions, such as nutrient limitations. The stringent response is a near-ubiquitous bacterial stress response which is mediated by accumulation of two hyper-phosphorylated derivatives of GTP and GDP: guanosine pentaphosphate (pppGpp) and guanosine tetraphosphate (ppGpp), collectively referred to as (p)ppGpp (Hauryliuk et al., 2015). The stringent response and (p)ppGpp have important roles in the regulation of bacterial virulence, survival during host invasion, and antibiotic resistance (Dalebroux et al., 2010;Dalebroux and Swanson, 2012;Hauryliuk et al., 2015).

RelA/SpoT homolog (RSH) are ubiquitous bacterial enzymes responsible for synthesis and degradation of (p)ppGpp (Atkinson et al., 2011;Jimmy et al., 2019). In the majority of beta- and gamma-proteobacteria, including *Escherichia coli*, the RSHs are represented by two multi-domain enzymes, the namesakes of the protein family – RelA and SpoT. The ‘long’ multi-domain RSHs are comprised of two functional regions: the catalytic N-terminal (NTD) and the regulatory C-terminal (CTD) (Figure 1A) (Atkinson et al., 2011). The NTD encompasses the (p)ppGpp hydrolase domain (HD; enzymatically inactive in RelA) as well as a (p)ppGpp synthetase domain (SYNTH). The CTD is comprised of four domains: Thr-tRNA synthetase, GTPase and SpoT domain (TGS domain); helical domain; zing finger domain (ZFD); and the RNA recognition motif (RRM). RelA is the best studied of all of the RSH representatives and its regulation is the focus of this study.

**Figure 1.**
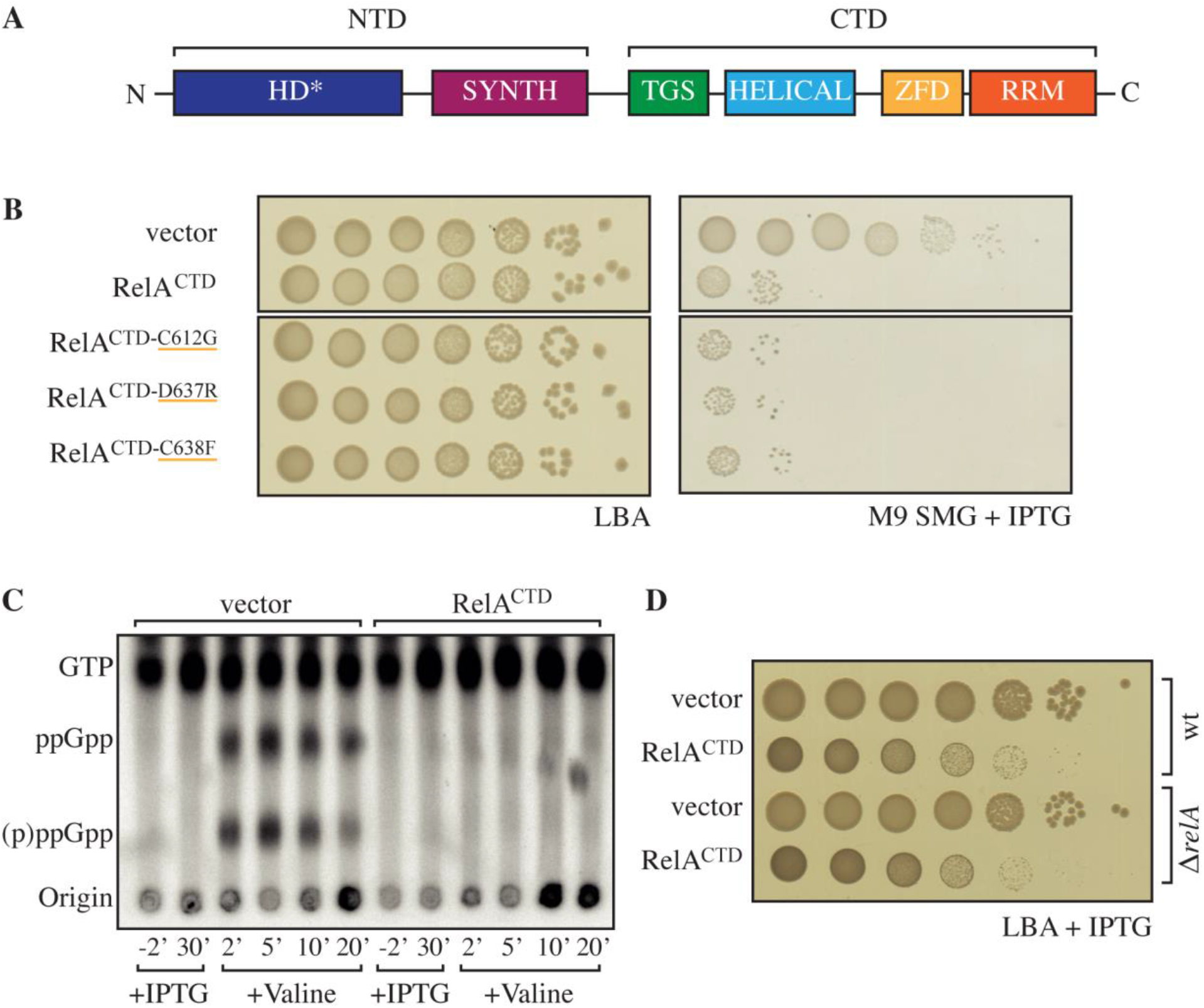
High-level ectopic expression of RelA^CTD^ is inhibitory to growth independently of native RelA’s activity. **(A)** Domain structure of RelA, the enzymatically-inactive (p)ppGpp hydrolysis (HD*) and a functional (p)ppGpp synthesis (SYNTH) NTD domains. TGS (ThrRS, GTPase and SpoT), Helical, ZFD (Zinc Finger Domain) and RRM (RNA recognition motif) domains comprise the regulatory CTD region. **(B)** *E. coli* MG1655 (wt) cells were transformed with high copy IPTG inducible vector, pMG25 (vector), pMG25::*relA^CTD^*, pMG25::*RelA^CTD-C612G^*, pMG25::*RelA^CTD-D637R^*, and pMG25::*RelA^CTD-C638F^*. The transformed cells were grown at 37 °C overnight in M9 minimal medium with 100 μg/ml ampicillin. Ten-fold serial dilutions were made and spotted onto LB agar (LBA) supplemented with 100 μg/ml ampicillin, as a loading control, and onto SMG plates supplemented with 100 μg/ml ampicillin and 1 mM IPTG to induce *relA* variant expression. **(C)** Representative audioradiogram of a PEI Cellulose TLC plate showing (p)ppGpp accumulation of *E. coli* MG1655 carrying pMG25 (vector) or pMG25::*relA^CTD^* upon valine-induced isoleucine starvation. See Materials and Methods for more details. **(D)** Overnight cultures of *E. coli* MG1655 (wt) and MG1655Δ*relA*(Δ*relA*) transformed with pMG25 (vector) or pMG25::*relA^CTD^*, grown in LB with 100 μg/ml ampicillin. Ten-fold serial dilutions of were made and spotted onto LB agar supplemented with 1 mM IPTG.

While RelA has no detectable hydrolytic activity (Shyp et al., 2012), it possesses a strong ribosome-dependent (p)ppGpp synthetic activity that is activated by deacylated tRNA in the ribosomal A-site – ribosomal ‘starved’ complexes (Haseltine and Block, 1973). In the absence of the starved ribosomal complexes, RelA displays a very low (p)ppGpp synthetic activity that is not stimulated by deacylated tRNA (Wendrich et al., 2002;Knutsson Jenvert and Holmberg Schiavone, 2005;Kudrin et al., 2018). Deletion of the CTD results in (p)ppGpp accumulation in the cell (Schreiber et al., 1991;Svitil et al., 1993;Gropp et al., 2001), suggesting that the CTD is essential for regulation *via* repression of the synthetic activity of RelA.

Two opposing models have been presented for the regulatory role of the CTD. The first was proposed by three independent groups (Gropp et al., 2001;Yang and Ishiguro, 2001;Jain et al., 2006) suggesting that under optimal growth conditions, RelA exists as a synthetically inactive dimer and upon amino acid starvation the enzyme is activated in the monomeric form. The dimerization was proposed to be mediated by the Zinc Finger Domain (ZDF; alternatively referred to as CC, conserved cysteine as per (Atkinson et al., 2011)), and mutations C612G, D637R, and C638F were proposed to abrogate dimerization (Gropp et al., 2001;Jain et al., 2006). Importantly, the ZDF is crucial for RelA interaction with ribosome, forming specific contacts with the A-site Finger rRNA element that acts as an anchoring point for RelA:ribosome complex formation (Arenz et al., 2016;Brown et al., 2016;Loveland et al., 2016;Kudrin et al., 2018). The second model was proposed by a study of a bi-functional ‘long’ RSH enzyme Rel from *Streptococcus equisimilis* (Rel_*Seq*_) which suggested that the CTD of Rel represses the (p)ppGpp synthesis by NTD *via* intramolecular contacts *in cis* (Mechold et al., 2002).

Here, we provide evidence that *E. coli* RelA is predominantly regulated through intramolecular (autoinhibition of the NTD by the CTD *in cis*) rather than intermolecular (dimerization, autoinhibition *in trans*) interactions.

## 2 Materials and Methods

### 2.1 Bacterial strains, plasmids and growth conditions

The strains and plasmids used in this study are listed in Table 1. Oligonucleotides used in this study are provided in Supplementary Table 1. For growth assays in Luria–Bertani (LB, BD Difco – Fisher Scientific) medium or on LB agar, the relevant strains were grown overnight in LB medium supplemented with antibiotic selection. For growth assays on SMG (Serine, Methionine and Glycine) plates (Uzan and Danchin, 1978), the strains were grown overnight in M9-glucose minimal medium (1 x M9 salts, 1 μg μl^−1^ thiamine, 1 mM MgCl_2_, 0.1 mM CaCl, pH 7.4; 10 x M9 salts: 80 g Na_2_HPO_4_, 30 g KH_2_PO_4_, 5 g NaCl per 1 L) supplemented with antibiotics for plasmid selection. The SMG plates were made by supplementing M9-glucose minimal medium with 0.7% agarose, 100 μg/ml of Serine, Methionine and Glycine and appropriate antibiotic selection. Expression from LacI-regulated promoters on solid or in liquid media was induced by the addition of 1 mM of IPTG. Bacterial cultures were routinely grown overnight at 37 °C, and plates were incubated at 37 °C for 18 h. 30 μg/ml ampicillin was used to select for low copy number plasmid pNDM220 and its derivatives, and 100 μg/ml ampicillin was used to select for the high copy number plasmid pMG25 and its derivatives. Growth assays present in figure panels originate from a single plate that were brought together, where applicable, this is indicated by a gap. All growth curves were performed in flasks, grown at 37 °C, with shaking at 160 rpm, unless otherwise indicated.

**Table 1.**
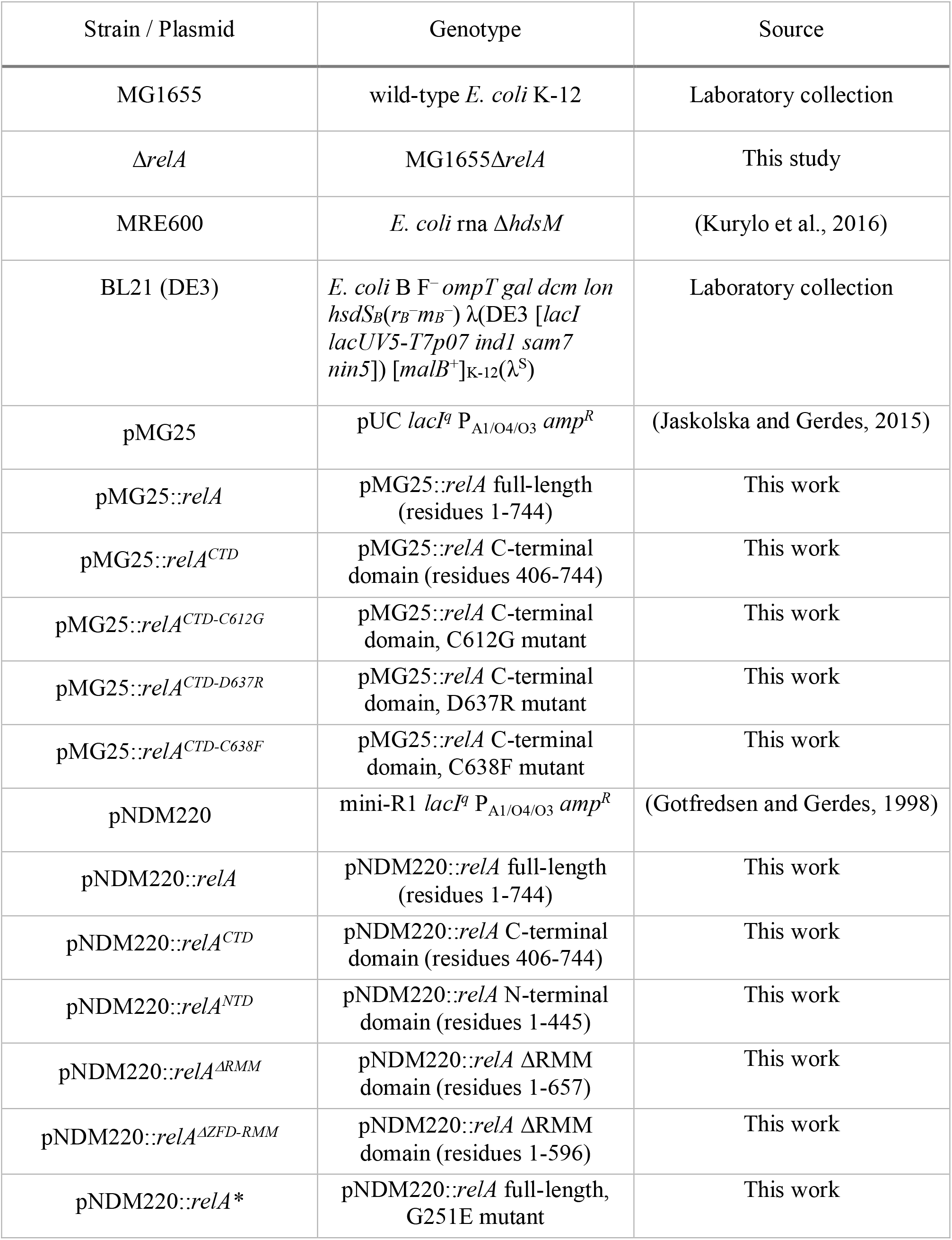
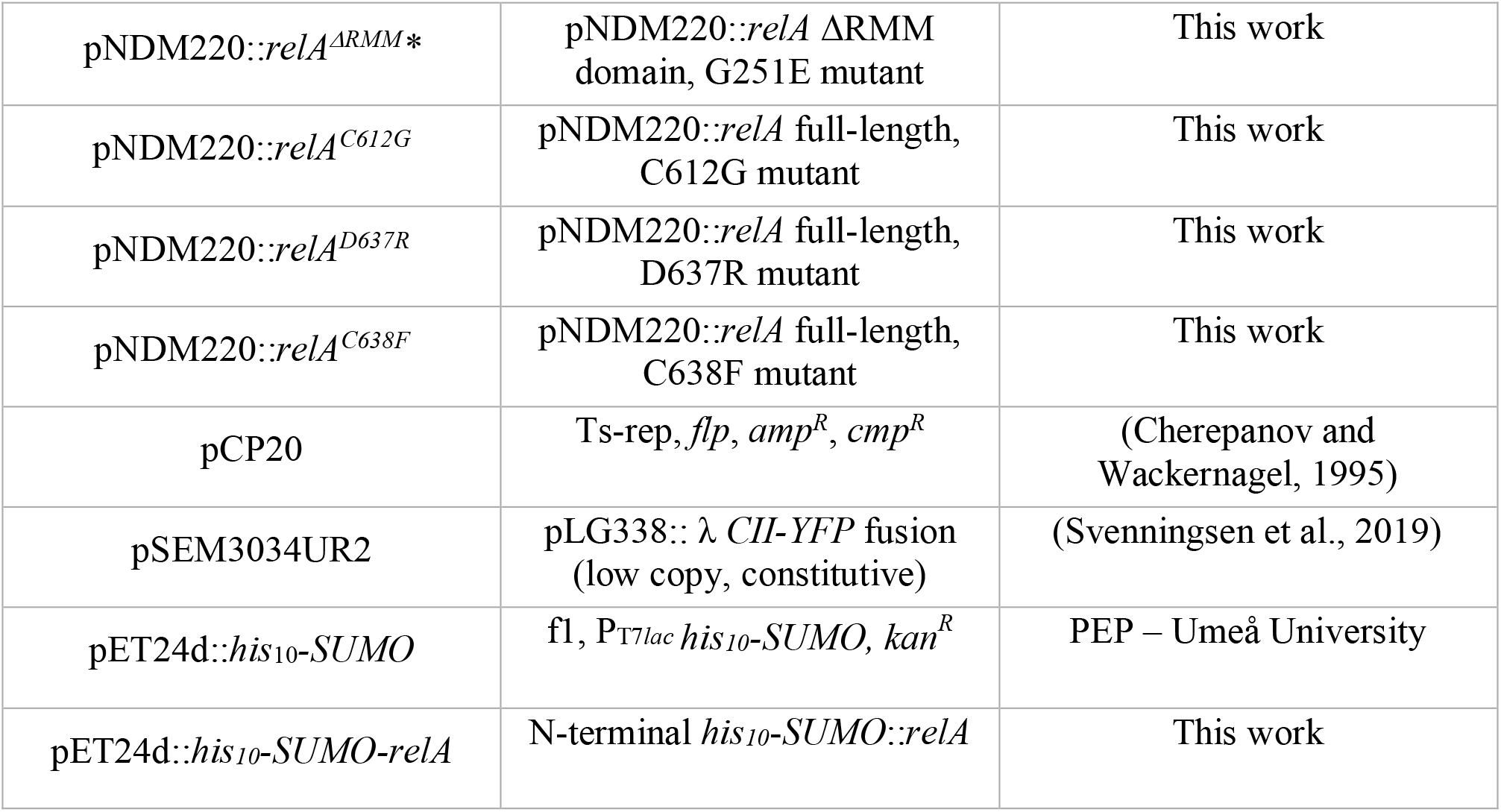
Bacterial strains and plasmids used in this study.

### 2.2 (p)ppGpp measurements by Thin Layer Chromatography (TLC)

To determine the (p)ppGpp levels in cells we modified the method from (Cashel, 1994;Tian et al., 2016). In brief, overnight cultures were diluted 100-fold in 5 ml of MOPS glucose minimal medium supplemented with all five nucleobases (10 μg/ml of each, Sigma Alderich) and incubated at 37 °C with shaking. All of the strains had similar growth rates in this medium. At an optical density at 600 nm (OD_600_) of ≈0.5, cells were diluted 10-fold to an OD_600_ of ≈0.05 and were left to grow with shaking at 37 °C with H_3_^32^PO_4_ (100 μCi/ml). When looking at the effect of RelA^CTD^ expressed from the low copy number plasmid (Figure 2C and Supplementary Figure 4), 1 mM IPTG was also added at this point. The cells were then grown for ≈2 generations (OD_600_ ≈0.2) before amino acid starvation was induced by the addition of either 500 μg/ml valine (Sigma Aldrich) or 400 μg/ml serine hydroxamate (Sigma Aldrich). Whereas, due to its inhibitory effect on growth, RelA^CTD^ was expressed from the high copy number plasmid for 30 min after the cells were grown for ≈2 generations (OD_600_ ≈0.2). Subsequently, amino acid starvation was induced by the addition of valine (Figure 1C). Fifty-microliter samples were withdrawn before and at various times (see figures) after the induction of amino acid starvation. With the induction of RelA and RelA^ΔRRM^ (Figure 3C), after labelling of cells for ≈2 generations, 1 mM IPTG was added and 50 μl samples were taken at −2, 10, 30, and 60 min. The reactions were stopped by the addition of 10 μl of ice-cold 2 M formic acid. A 10 μl aliquot of each reaction mixture was loaded on polyethyleneimine (PEI) cellulose thin-layer chromatography (TLC) plates (GE Healthcare) and separated by chromatography in 1.5 M potassium phosphate at pH 3.4. The TLC plates were revealed by phosphorimaging (GE Healthcare) and analysed using ImageQuant software (GE Healthcare). The increase in the level of (p)ppGpp was normalized to the basal level (time zero) for each strain.

**Figure 2.**
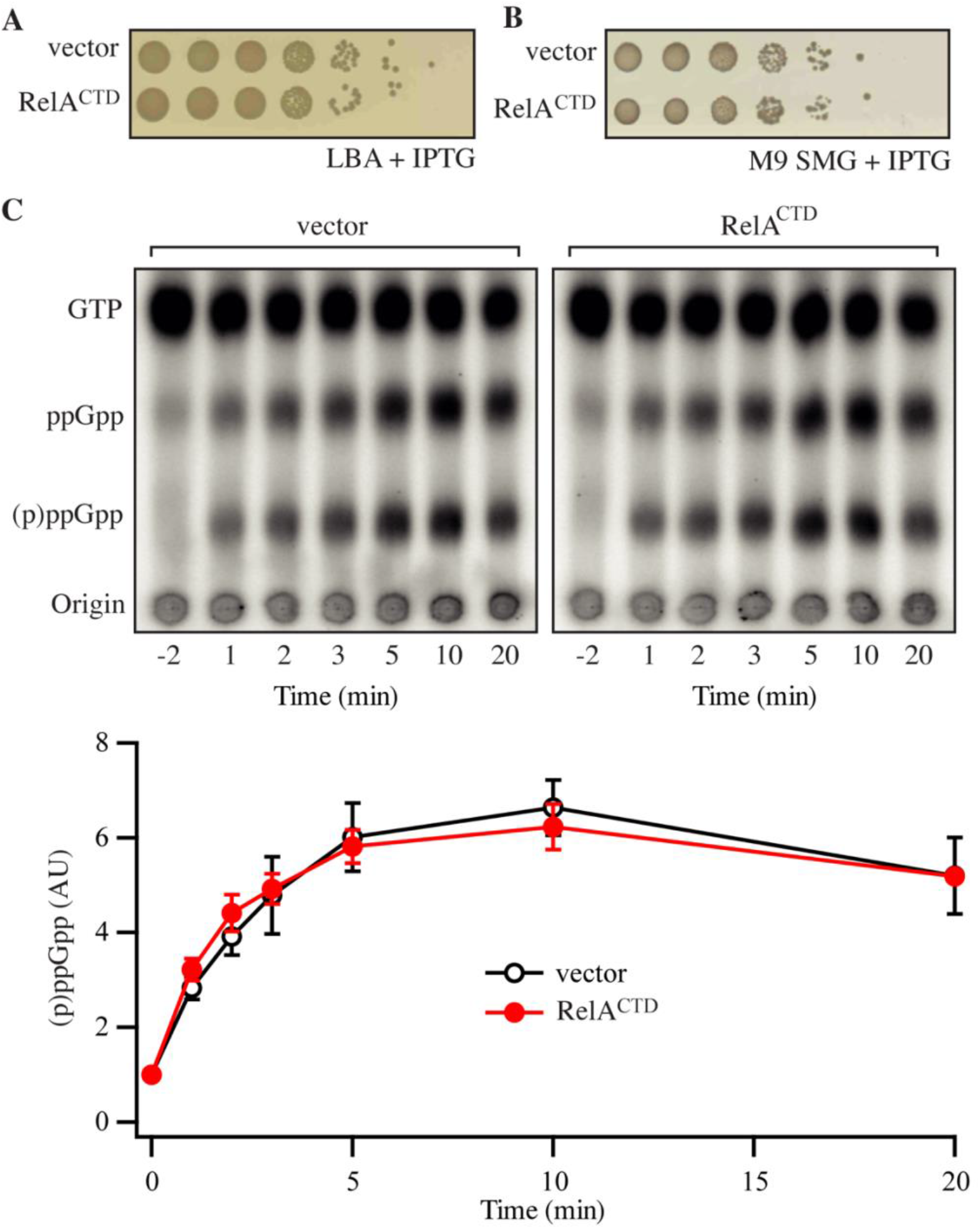
Low-level ectopic expression of RelA^CTD^ does not interfere with the functionality of endogenous RelA. **(A)** and **(B)** Overnight cultures of *E. coli* MG1655, transformed with low copy IPTG inducible vector pNDM220 (vector) or pNDM220::*relA^CTD^*, were grown at 37 °C overnight in M9 minimal medium with 30 μg/ml ampicillin. Ten-fold serial dilutions were made and spotted onto LB agar (LBA) and SMG plates, both supplemented with 30 μg/ml ampicillin and 1 mM IPTG. **(C)** Representative audioradiogram of a PEI Cellulose TLC plate showing (p)ppGpp accumulation of *E. coli* MG1655 carrying pNDM220 (vector) or pNDM220::*relA^CTD^* upon valine-induced isoleucine starvation. The curves represent the average fold increase for three independent measurements, and the error bars represent standard errors. The levels of (p)ppGpp were normalized to the pre-starved level for each strain at −2 minutes. See experimental procedures for more details.

### 2.3 Monitoring translation efficiency

Overnight *E. coli* MG1655 cells harbouring the CII-YFP plasmid pSEM3034UR2 kindly donated by Szabolcs Semsey and pMG25 (−ve), pMG25::*relA*, pMG25::*relA^G251E^*, pMG25::*relA^CTD^* were grown into balanced growth in MOPS glucose minimal medium, before back-dilution to 0.05 OD_600_ into flat-bottom 96-well microtiter plate (Sterilin, Thermo Scientific) and placed in Synergy H1 (BioTek) microplate reader. Subsequently, the cultures were grown at 37 °C with continuous orbital shaking. Every 15 minutes growth measured by taking the optical density (OD_600_), and the fluorescence was read from the bottom at 485 nm excitation and the 520 nm emission wavelengths (Figure 3).

**Figure 3.**
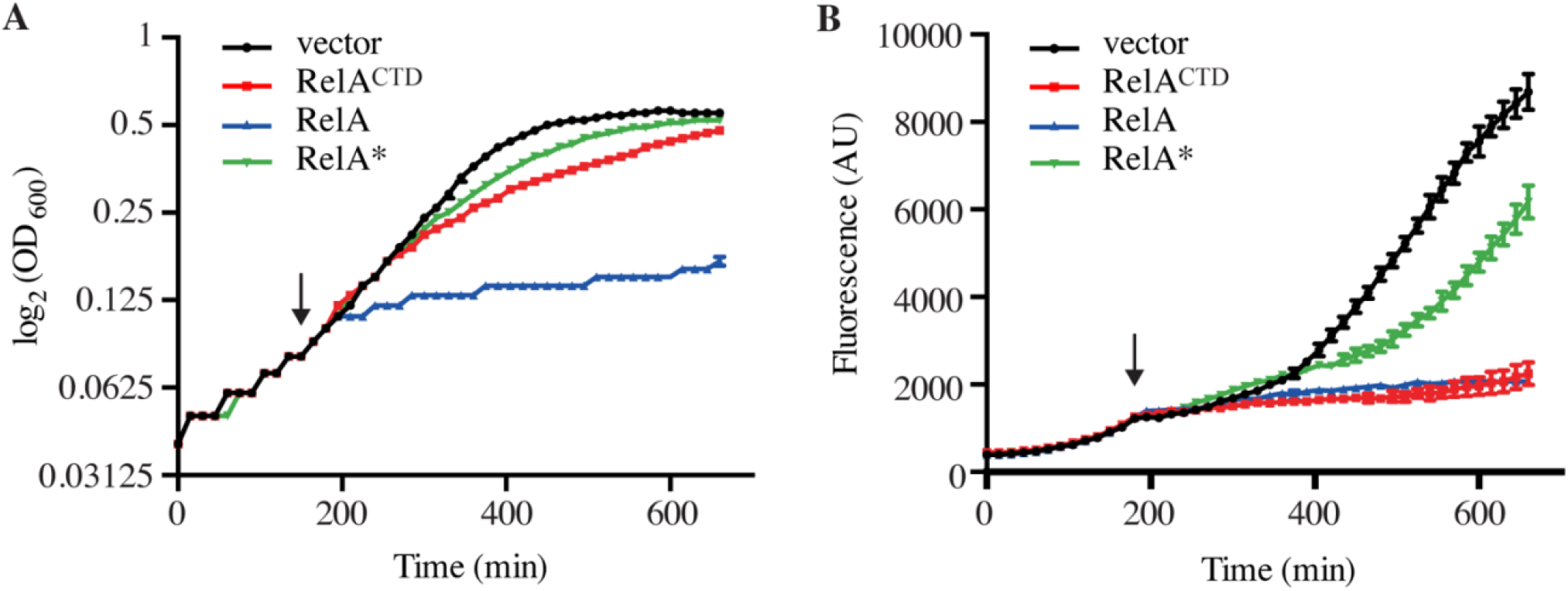
High-level ectopic expression of RelA^CTD^ impairs translation. Exponentially growing *E. coli* MG1655 cells harbouring the CII-YFP plasmid were back-diluted to OD_600_ 0.05, before the expression of *relA*, catalytically-inactive G251E mutant *relA* *, or *relA^CTD^* from high copy vector pMG25 was induced by addition of 1 mM IPTG (indicated by the arrow). **(A)** Growth, measured at 600 nm (OD_600_) and **(B)** the fluorescence of YFP, measured at 520 nm was monitored every 15 minutes for 11 h. See materials and methods for more details.

### 2.4 Western blotting

For the detection of overproduced RelA and RelA^ΔRRM^ the appropriate plasmid was induced as described in the text and samples taken at the indicated time points (Figure 4). At each time point, 1ml samples were taken and the cells were harvested by centrifugation at 4 °C. Subsequently, the cell pellet was resuspended in an appropriate volume of 1x LDS loading buffer (Life Technologies) and boiled for 5 min. 20 μl of the samples was then resolved on a 4-12% NuPAGE gel (Life Technologies). Subsequently, the proteins were transferred using a semi-dry blotting apparatus (Hoefer Scientific Instruments) to a PVDF membrane (Amersham) and incubated with the primary antibody in PBS with 0.1% Tween-20 and 5% milk powder. The primary antibodies used in this study were raised in rabbit against a peptide sequence from the ZFD of RelA (Eurogentec). Next, membranes were incubated with HR conjugated anti-rabbit IgG antibody (Sigma) in PBS with 0.1% Tween-20 and 5% milk powder. Finally, the protein bands were visualised using Pierce ECL chemiluminescence substrate (Thermo Scientific) according to manufacturer’s instructions and the signal was detected in an Imagequant LAS4100.

**Figure 4.**
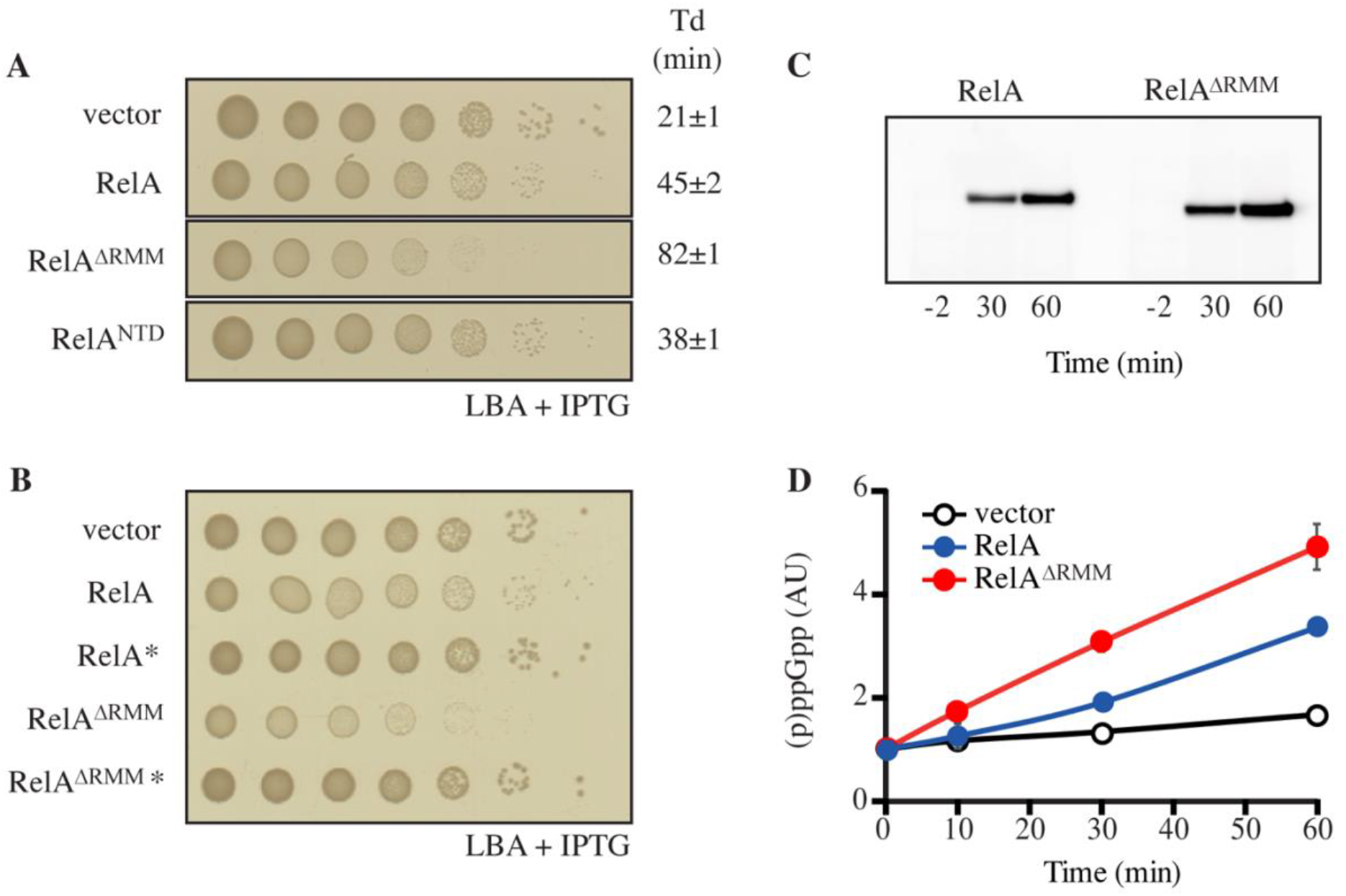
Truncation of RelA leads to accumulation of (p)ppGpp. **(A)** and **(B)** *E. coli* MG1655 cells were transformed with low copy IPTG inducible vector, pNDM220 (vector), pNDM220::*relA*, pNDM220::*relA*\*, pNDM220::*relA^ΔRRM^*, pNDM220::*relA^ΔRRM^*\*, or pNDM220::*relA^NTD^*. Ten-fold serial dilutions of overnight LB cultures were made and spotted onto LB agar (LBA) supplemented with 30 μg/ml ampicillin and 1 mM IPTG. Loading controls are presented in Supplementary Figure 6B. **(C)** *E. coli* MG1655Δ*relA* transformed with pNDM220::*relA* or pNDM220::*relA^ΔRRM^* were grown at 37 °C to OD_600_ 0.5, before 1 mM IPTG was added for plasmid induction. Samples were withdrawn at the mentioned times (minutes after induction) and western blotting was performed as described in the experimental procedures. **(D)** *E. coli* MG1655 carrying pNDM220 (vector), pNDM220::relA, or pNDM220::*relA^ΔRRM^* were grown exponentially in MOPS minimal medium with 30 μg/ml ampicillin before1 mM IPTG was added, for plasmid induction at time 0 minutes. The curves represent the average fold increase of (p)ppGpp for two independent measurements and the error bars represent standard errors. The levels of (p)ppGpp were normalized to the pre-starved level (−2 minutes) for each strain.

### 2.5 Purification of *E. coli* 70S ribosomes and untagged native RelA

*E. coli* 70S ribosomes were prepared from RNase I-deficient *E. coli* strain MRE600 (Kurylo et al., 2016). Bacteria were grown in 2YT medium (Sigma Alderich) to OD_600_ ≈ 0.5, collected by centrifugation, and the ribosomes were purified by sucrose gradient centrifugation as described for preparation of *Enterococcus faecalis* 70S earlier (Murina et al., 2018).

For purification of RelA, *E. coli* BL21 DE3 harbouring pET24d::*his_10_-SUMO-relA* expression construct were grown, induced, harvested and lysed as previously described (Kudrin et al., 2018). All liquid chromatography steps were performed using ÄKTA Avant 25 system and chromatographic columns from GE Healthcare were used. In order to exclude a possibility of substitution of Zn^2+^ ions in RelA’s ZFD for Ni^2^+ during purification on metal affinity chromatography column (Block et al., 2009), HisTrap 5 HP column was stripped from Ni^2+^ (according to manufacturer recommendations) and loaded with 10 ml of 100 mM Zn(OAc)_2_, pH 5.0. The column was then washed with four column volumes of deionized water and pre-equilibrated with four column volumes of binding buffer (25 mM Hepes pH 7.6, 320 mM NaCl, 10 mM imidazole, 5 mM MgCl_2_, 4 mM BME, 10% glycerol). Clarified cell lysate (≈50 ml) was applied on the column at the flow rate 5 ml/minute. Then the column was washed with binding buffer (2.5 column volumes) and the protein was eluted with six column volumes of 0-100% gradient of elution buffer (binding buffer with 500 mM imidazole) and 2 ml fractions were collected into 96 deep well plates (Omega Bio-tek). The collected fractions were run on SDS-PAGE gel and the fractions corresponding to His_10_-SUMO-RelA with the least nucleic acid contamination was collected (≈5 ml, highlighted in Figure 5A; corresponding SDS PAGE analysis is shown on Figure 5D). These fractions were applied to Hiprep 10/26 desalting column pre-equilibrated with the storage buffer (25 mM Hepes pH 7.6, 720 mM KCl, 50 mM arginine, 50 mM glutamic acid, 10 mM imidazole, 5 mM MgCl_2_, 4 mM BME, 10% glycerol) (Figure 5B). The peak was collected and concentrated to the volume not bigger than 5 ml using Amicon centrifugal filters with 50 kDa cut-off. His_6_-UlpI SUMO protease (Protein Expertise Platform (PEP), Umeå University) was added to the protein solution to a final concentration of 1.3 μM and incubated for 15 minutes at room temperature in order to cleave off His_10_-SUMO tag and the protein solution was applied on HisTrap 1 HP (GE Healthcare) loaded with Zn^2+^ and pre-equilibrated with storage buffer. The flow-through was collected and concentrated using Amicon concentrators with 50 kDa cut-off (Figure 5C). The concentrated protein was aliquoted per 20-30 μl, flash frozen in liquid nitrogen and stored at −80°C. The purity of RelA preparations was assessed by SDS-PAGE (Figure 5D) and spectrophotometrically (OD_260_/OD_280_ ratio below 0.8 corresponding to less than 5% RNA contamination (Layne, 1957)).

**Figure 5.**
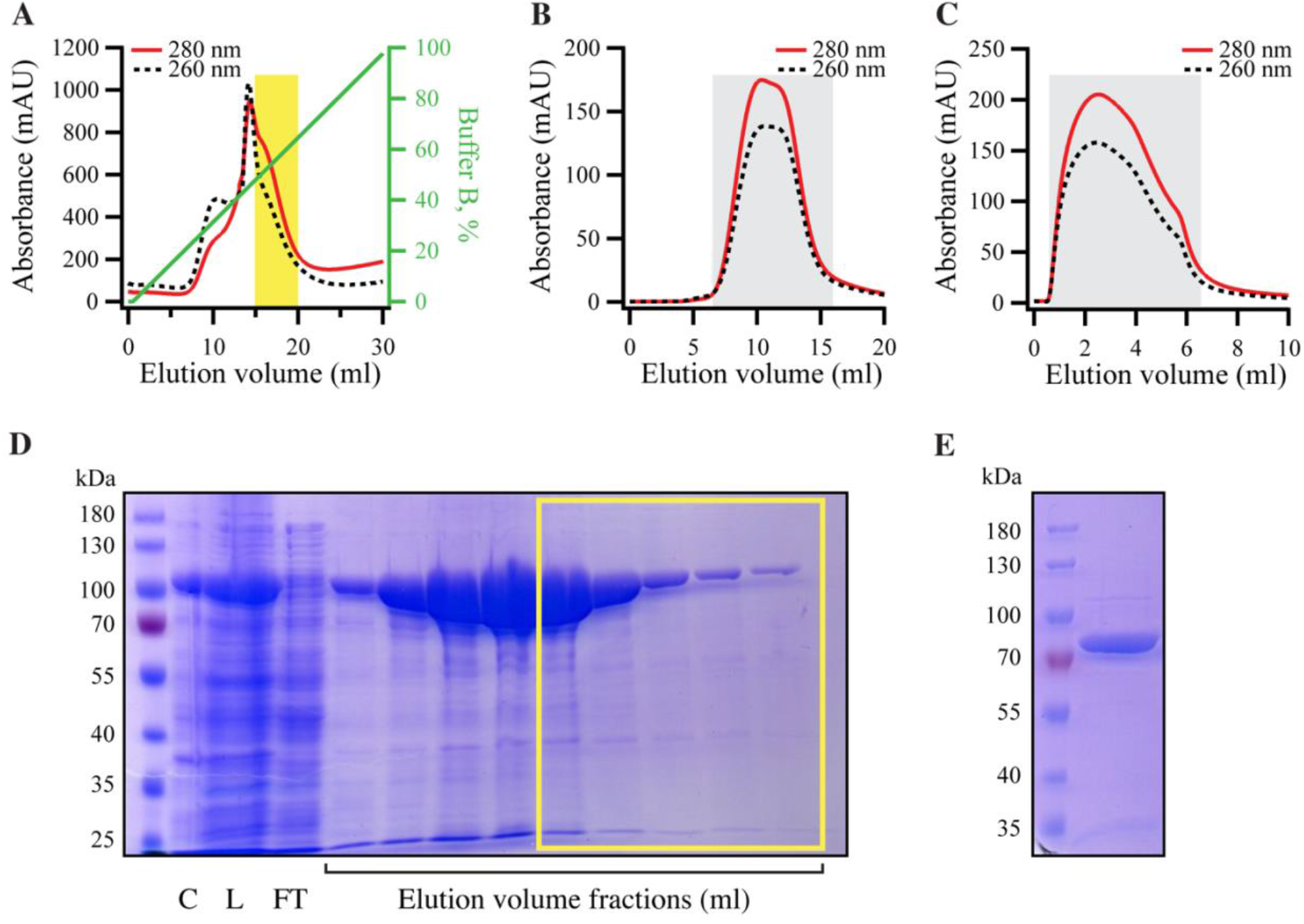
Purification of RNA-free untagged full-length *E. coli* RelA. N-terminally His_10_-SUMO tagged RelA was overexpressed and purified as described in detail in Materials and Methods. **(A)** Cells were lysed and subjected to immobilized metal affinity chromatography (IMAC). The fraction corresponding the RelA with the lowest contamination of nucleic acids (highlighted in yellow) was carried forward. **(B)** Following the buffer exchange, the greyed-out fractions were pooled and the His_10_-SUMO tag was cleaved off by the addition of His_6_-Ulp1. **(C)** His_6_-Ulp1 and the His_10_-SUMO tag was then purified from RelA by a second IMAC round. The greyed-out fractions were pulled, concentrated, aliquoted and stored −80 °C. **(D)** SDS-PAGE showing the eluted fractions from the first IMAC round **(A)**, fractions carried forward are highlighted with the yellow border. **(E)** SDS-PAGE analysis of purified native untagged RelA (≈84 kDa).

### 2.6 ppGpp synthesis assay

All experiments were performed at 37 °C in HEPES:Polymix buffer, pH 7.6, with 5 mM Mg^2+^ (Antoun et al., 2004). Since increased concentration of KCl inhibits RelA activity, RelA stock solution containing 720 mM KCl was first diluted 7.2 times with HEPES:Polymix buffer without KCl to final KCl concentration of 100 mM and then added to the reaction mixture. The reaction mixture containing 1 mM [^3^H]GDP (Perkin Elmer), 100 μM pppGpp as well as RelA at increasing concentrations (25, 50, 100, 200 and 400 nM), either supplemented or not by 0.6 μM 70S ribosomes, was preincubated for 3 minutes at 37 °C and then the reaction was started by adding pre-warmed ATP to a final concentration of 1.5 mM. During the time course of the reaction 5 μl aliquots of the reaction mixture was quenched with 4 μl of 70% formic acid supplemented with 4 mM GTP and GDP (for UV shadowing). Time points were centrifuged for 5 minutes at 14,000 rpm, the supernatant was resolved on PEI-TLC (Macherey-Nagel) in 0.5 KH_2_PO_4_ buffer (pH 3.5) and the spots corresponding to [^3^H]GDP and [^3^H]ppGpp were cut out and quantified by scintillation in 5 ml of ScintiSafe scintillation cocktail (Fisher Scientific).

## 3 Results

### 3.1 High-level ectopic expression of Rel^CTD^ is inhibitory to growth independently of endogenous RelA

To test the functionality of RelA-mediated stringent response we utilised the so-called SMG test (minimal medium plates supplemented with serine, methionine, and glycine plates). Growth of *E. coli* on SMG is strictly dependent on the functionality of RelA (Uzan and Danchin, 1978). To investigate the effect of high-level ectopic expression of RelA^CTD^ (pUC derivative pMG25::RelA^CTD^, P_A1/O4/O3_ promoter) on RelA activity, we assayed the growth of wild-type *E. coli* MG1655 (wt) cells growing on SMG. In agreement with the previous studies (Gropp et al., 2001;Yang and Ishiguro, 2001), high expression levels of RelA^CTD^ severely inhibit growth on SMG as compared to the empty vector control (Figure 1B). When *E. coli* is challenged with amino acid starvation, the levels of (p)ppGpp increase (Figure 1C, left side). At the same time, expression of RelA^CTD^ from the high-copy vector abrogates the response of native RelA to amino acid starvation and the levels of (p)ppGpp in the cells do not increase beyond the basal levels (Figure 1C, right side), i.e. native RelA fails to be activated in cells expressing high levels of RelA^CTD^. The lack of activation of native RelA by amino acid starvation upon high-level expression of RelA^CTD^ could be explained either *via* direct inhibition of full-length RelA by RelA^CTD^ (autoregulation *in trans*) or interference with the activation of the full-length RelA by starved ribosomal complexes.

Next, we tested the ZFD domain mutations C612G, D637R, and C638F that were previously shown to render RelA^CTD^ unable to suppress the activity of native RelA (Gropp et al., 2001). We individually introduced these three mutations into the high copy plasmid encoding RelA^CTD^ to see whether the negative dominance of the CTD is abolished. The mentioned amino acid changes do not affect the expression or the stability of C-terminal fragments (Gropp et al., 2001). To our surprise, in our hands, overexpression of mutant variants of RelA^CTD^ still inhibits the growth of wt cells on SMG (Figure 1B). Therefore, high levels of RelA^CTD^, with or without the mutated conserved residues, impair the functionality of RelA-mediated stringent response.

To test whether the observed effect of RelA^CTD^ overexpression on the stringent response is solely mediated by inactivation of endogenous RelA or *via* other mechanism(s) of growth inhibition, we investigated the effect of RelA^CTD^ expression in wt as well as Δ*relA* backgrounds in both rich liquid and solid medium under unstarved conditions where functional RelA is not required. Surprisingly, upon induction of RelA^CTD^, the growth rate is reduced identically in the wt and Δ*relA* backgrounds, both in liquid and solid media indicative of the RelA-independent mechanism of growth inhibition (Figure 1D, Supplementary Figure 1A). Additionally, high expression levels of RelA^CTD^ bearing C612G, D637R, and C638F mutations are also inhibitory to growth in unstarved conditions (Supplementary Figure 2A), further reinforcing the idea growth inhibition being independent of the functionality of the endogenous RelA and the stringent response.

### 3.2 Low-level ectopic expression of Rel^CTD^ does not affect the activation of endogenous full-length RelA

Since high-level ectopic expression of RelA^CTD^ was inhibitory to growth independently of endogenous RelA (Figure 1D), no conclusions could be drawn with regard to its effect on the regulation of RelA. Ectopic expression of RelA^CTD^ from a high copy number plasmid results in accumulation of high levels of the recombinant protein in the cell (Supplementary Figure 1B). Given that there are only a few hundred molecules of RelA in the cell (Pedersen and Kjeldgaard, 1977;Justesen et al., 1986;Li et al., 2014;Schmidt et al., 2016), we switched to a very low copy number mini R1 plasmid (pNDM220, 1-2 copy per chromosome (Molin et al., 1979); P_A1/O4/O3_ promoter). We tested whether expression of the full-length RelA from this plasmid (pNDM220::*relA*) can restore the growth of a strain lacking *relA* (*E. coli* MG1655Δ*relA*) on SMG. As seen from Supplementary Figure 3, expression of RelA in a Δ*relA* background from this low copy number plasmid supports growth similar to wt cells. Notably, RelA bearing C612G, D637R, and C638F mutations in the ZFD domain also show full complementation (Supplementary Figure 3). As expected, the empty plasmid control and the catalytically-inactive RelA mutant G251E (commonly designated as RelA* (Gropp et al., 2001)) fails to grow on SMG (Supplementary Figure 3). This result indicates that the expression from this plasmid resulted in more physiologically-relevant levels of RelA.

Next, we repeated the experiments presented in the section 3.1 using ectopic expression of RelA^CTD^ from the low copy number R1 plasmid (pNDM220::*relA^CTD^*, P_A1/O4/O3_ promoter). As seen from Figure 2A, lower levels of RelA^CTD^ are not inhibitory to growth under unstarved conditions on LB agar. Likewise, at such expression levels, RelA^CTD^ does not affect the growth of starved wt cells on SMG plates (Figure 2B) or the out growth in SMG media (Supplementary Figure 5B), suggesting that native RelA in these conditions is still active. We then tested the profile of the of RelA mediated response upon amino acid starvation by measuring (p)ppGpp produced in the cell. As evident from the SMG plate assays (Figure 2B), low levels of RelA^CTD^ do not abrogate the functionality of endogenous RelA upon induction of the stringent response by addition of either valine that causes amino acid starvation in K12-based *E. coli* strains (Leavitt and Umbarger, 1962) (Figure 2C) or serine hydroxamate (SHX), the competitive inhibitor of seryl-tRNA synthetase (Tosa and Pizer, 1971) (Supplementary Figure 4). Furthermore, in case additional domains other than the CTD are required, we wanted to explore the possibility of dimerization of full-length RelA enzyme. We hypothesised that if RelA is capable of dimerization, that endogenous expression of a catalytically inactive RelA mutant would bind and titrate out the native, catalytically active RelA, shifting the equilibrium away from free and active RelA in the cell, thus impairing the response to amino acid starvation. In order to test this theory, we repeated the above experiments with the inactive RelA G251E mutant, RelA* (pNDM220::*relA*\*). As seen from Supplementary Figure 5, low-level expression of RelA* does not affect growth on SMG plate or media, and does not affect the activation of endogenous RelA upon induction of amino acid starvation. Taken together, our data indicate that the activation of RelA is not predominantly regulated through intermolecular interactions.

### 3.3 High-level ectopic expression of RelA^CTD^ impairs translation

The above data obtained using low expression vector suggests that RelA^CTD^ does not inhibit RelA activation by forming dimers. However, the question remained as to why high levels of RelA^CTD^ are inhibitory to growth (Figure 1D). All recent studies have unanimously showed that the CTD is critical for RelA binding to the ribosome (Agirrezabala et al., 2013;Arenz et al., 2016;Brown et al., 2016;Loveland et al., 2016;Winther et al., 2018). Furthermore, biochemical studies have shown that upon overexpression of RelA^CTD^, the bulk of the protein is ribosome bound (Yang and Ishiguro, 2001). Therefore, we hypothesised that the reduced growth rate associated with high levels of RelA^CTD^ could be due to impairment of translation. To monitor the efficiency of translation upon expression of RelA variants (RelA, RelA* and RelA^CTD^), we transformed wt cells with a low copy number plasmid encoding bacteriophage λ CII protein fused to Venus yellow fluorescence protein (CII-YFP) which allows the indirect monitoring of translation via measuring of fluorescence (Svenningsen et al., 2019).

As expected, wt cells with the empty vector exhibited increased levels of fluorescence correlated with growth, demonstrating active translation (Figures 3A and B). Wild-type cells harbouring CII-YFP were grown exponentially before RelA and its variants were induced. High-level expression of full-length RelA results in accumulation of (p)ppGpp to levels at least as high as those seen during the stringent response (Schreiber et al., 1991;Svitil et al., 1993;Gropp et al., 2001). The alarmone is a potent inhibitor of bacterial growth targeting both transcription (Sorensen et al., 1994;Molodtsov et al., 2018), translation (Milon et al., 2006;Mitkevich et al., 2010) and ribosome assembly (Corrigan et al., 2016). Thus, as expected, upon over-expression of RelA, growth is immediately inhibited (Figure 3A). Consequently, the levels of fluorescence fail to increase, indicating arrest of translation (Figure 3B). Ectopic expression of RelA^CTD^ in mid-exponentially growing cells results in slower rather than total arrest of growth (Figure 3A), but the levels of fluorescence, as in the case of expression of the full-length RelA, do not increase suggesting translation is impaired (Figure 3B). Next, to test whether the presence of the NTD could regulate the inhibitory effect of the CTD, we checked the effect of expression of full-length catalytically inactive G251E mutant, RelA* (Supplementary Figure 3). As seen from Figure 3A, high-level expression of RelA* is inhibitory to growth, but to a lesser extent than RelA^CTD^. Moreover, the levels of fluorescence gradually increased upon expression of RelA* indicating continuous decrease of translational efficiency in these cells. Notably, the inhibitory effect of RelA^CTD^ and RelA* in the cell is more pronounced when overnight cultures were back diluted to a lower optical density and allowed more time to accumulate the expressed product, suggesting that the inhibition of growth is dose-dependent (Supplementary Figure 2B). Taken together, these results suggest that RelA^CTD^ is inhibitory to growth as translation is impaired upon expression. RelA* is less toxic to growth and the presence of the NTD appears to moderate the toxic effect of the CTD.

### 3.4 The RRM domain is crucial for regulation of RelA’s enzymatic activity

The data presented above are suggestive of RelA being primely regulated, not via dimerization (*in trans*), but intramolecularly (*in cis*). To probe the autoregulatory roles of the individual domains, we systematically removed domains of the regulatory CTD region and tested the effects of their expression on growth (Figure 4A, Supplementry Figure 6A). Low-level expression of full-length RelA from pNDM220 in wt cells, was slightly inhibitory to growth (Figure 4A). Compared to the empty vector control, expression of RelA yielded smaller colonies on solid media and a reduced growth rate in LB liquid cultures of 21 to 45 minutes (Figure 4A, Supplementry Figure 6C), presumably due to elevated levels of (p)ppGpp. The expression of the constitutively active, but unstable, RelA^NTD^ (Schreiber et al., 1991;Svitil et al., 1993;Gropp et al., 2001) from the same plasmid resulted in similar growth inhibition (Figure 4A, Supplementry Figure 6C). Surprisingly, expression of RelA lacking the RRM domain (RelA^ΔRRM^, Figure 1A), resulted in a more severe reduction of the growth rate than wild type RelA or the constitutively active RelA^NTD^, as observed by a comparatively smaller colony size and an increased doubling time of 86 minutes (Figure 4A, Supplementry Figure 6C). Western blot analysis showed that upon expression, RelA^ΔRRM^ has similar abundance as RelA in the cells (Figure 4C). Thus, showing that the RRM domain is essential for the regulation of the (p)ppGpp synthetic activity of RelA.

To test if the growth inhibition by RelA^ΔRRM^ is mediated via over-production of (p)ppGpp, we repeated the plate assay described above using RelA and RelA^ΔRRM^ variants harbouring the G251E mutation that abrogates RelA’s catalytic activity (Gropp et al., 2001). As seen in Figure 4B, the inhibition of growth is indeed dependent on the synthetic activity of RelA and RelA^ΔRRM^. To further substantiate this hypothesis, we directly measured the levels of (p)ppGpp upon expression of RelA and RelA^ΔRRM^ in exponentially growing wt cells. Expression of RelA resulted in a gradual increase in the (p)ppGpp levels over time, and, consistent with a stronger growth inhibition (compare Figures 4A and B, Supplementry Figure 6C) the accumulation of the alarmone was more pronounced in the case of RelA^ΔRRM^ (Figure 4D). Taken together, our results collectively suggest that the RRM domain is crucial for preventing uncontrolled (p)ppGpp production by RelA.

### 3.5 Synthetic activity of native untagged RelA is not inhibited by increasing enzyme concentrations

The concentration of RelA in *E. coli* is in the range of 50-100 nM (Pedersen and Kjeldgaard, 1977;Justesen et al., 1986;Li et al., 2014;Schmidt et al., 2016). We hypothesised that if there is regulation *via* dimerization is physiologically relevant, the dimerization affinity should be characterised by the equilibrium affinity constant (*K_D_*) in this concentration range, and, therefore, at the concentrations above the *K_D_* RelA would dimerize and become catalytically inhibited.

To test this hypothesis in the test tube, we purified native untagged *E. coli* RelA (Figure 5). Using untagged version of the protein for enzymatic studies is important since even addition of a C-terminal His_6_ tag can interfere with RelA’s regulation (Kudrin et al., 2018). To do so, we first subjected N-terminally His_10_-SUMO tagged RelA to immobilized metal affinity chromatography (IMAC). Since RelA makes extensive contacts with tRNA and ribosomal RNA which regulate its enzymatic activity (Arenz et al., 2016;Brown et al., 2016;Loveland et al., 2016;Kudrin et al., 2018;Winther et al., 2018), it was essential to purify the protein from nucleic acid contamination that is observed by monitoring both 260 nm and 280 nm UV traces during the IMAC purification step (Figure 5A). Therefore, we pooled only the fractions containing RelA with the lowest amounts of nucleic acid contamination, indicated by a yellow box in Figure 5A and D, rather than collecting the factions that contained the highest concentration of the protein as judged by absorbance at 280 nm and SDS PAGE analysis. Following a buffer exchange (Figure 5B), the His_10_-SUMO tag was cleaved by the addition of Ulp1, resulting in native untagged RelA (≈84 kDa), which was separated from the uncut version as well as the His_10_-SUMO tag by IMAC (Figure 5C). The purity of the protein preparation was checked by SDS-PAGE (Figure 5E), and importantly spectrophotometrically, giving the OD260/OD280 ratio of 0.8 corresponding to less than 5% RNA contamination (Layne, 1957).

We tested the synthetic activity of RelA in the range of concentrations from 25 to 400 nM, in the presence or absence of 0.6 μM 70S ribosomes (Figure 6). The enzymatic activity is largely insensitive to RelA concentration, suggesting that RelA is not regulated via through intermolecular interactions (dimerization).

**Figure 6.**
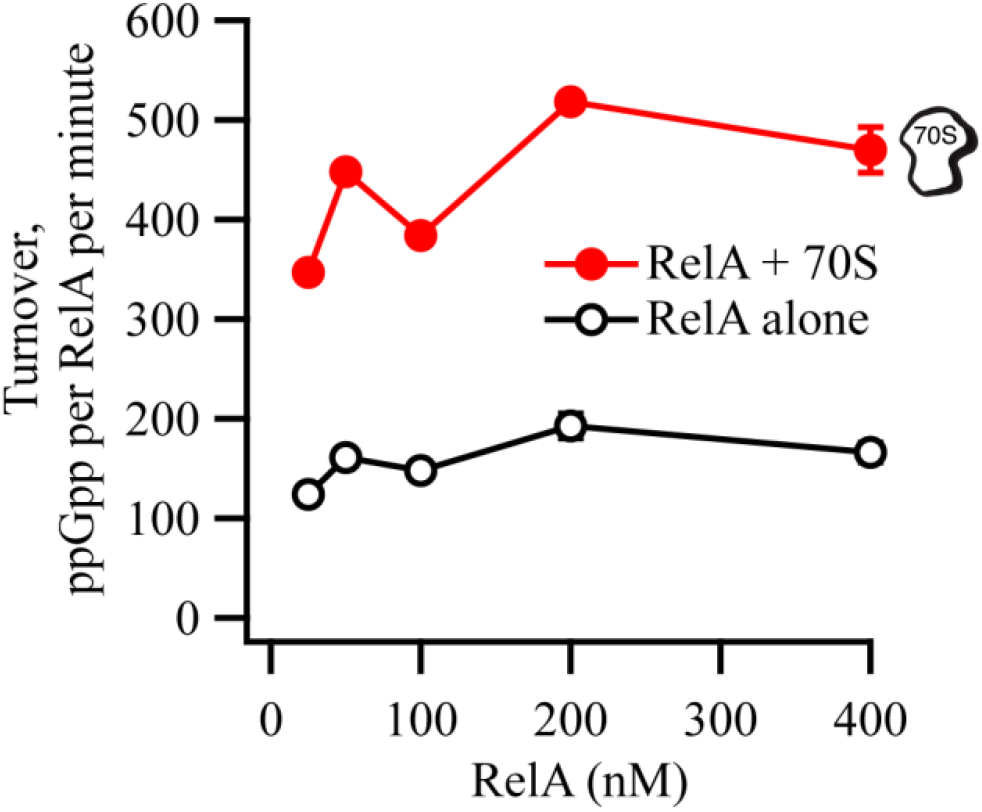
RelA’s ppGpp-synthetic activity is not inhibited by increasing concentrations of the enzyme. ppGpp synthetic activity of increasing concentrations of untagged native full-length RelA in the presence or absence of 0.6 μM *E. coli* 70S ribosomes, with 1 mM ^3^H-labelled GDP and 1.5 mM ATP. Enzymatic assays were performed at 37 °C in HEPES:Polymix buffer, pH 7.5, in the presence of 5 mM Mg^2+^. Error bars represent SDs of the turnover estimates by linear regression.

## 4 Discussion

In this report, we provide evidence that the catalytic activity of the *E. coli* stringent factor RelA, is unlikely to be regulated through oligomerization in the cell, as it was previously suggested (Gropp et al., 2001;Yang and Ishiguro, 2001;Jain et al., 2006). However, our results are consistent with an alternative model for RelA regulation via intradomain regulation *in cis*, proposed earlier for ‘long’ bifunctional RSH Rel from *Streptococcus equisimilis* (Mechold et al., 2002). This conclusion is substantiated by three lines of evidence.

The original ‘inhibition-via-dimerization’ model of RelA regulation, was motivated by the observation of high-level expression of RelA^CTD^ leading to inhibition of both growth and (p)ppGpp accumulation upon amino acid starvation (Gropp et al., 2001;Yang and Ishiguro, 2001). By performing additional control experiments overlooked in the original reports, we demonstrate that high-level expression of RelA^CTD^ is inhibitory to growth in a RelA-independent manner, likely *via* direct inhibition of translation. Recent studies provide a possible structural explanation of the microbiological results. The CTD region is responsible for anchoring RelA to the ribosome by forming multiple contacts with rRNA and A-site tRNA (Agirrezabala et al., 2013;Arenz et al., 2016;Brown et al., 2016;Loveland et al., 2016;Winther et al., 2018). Once bound to the ribosome, RelA^CTD^ would sterically clash with other ribosomal ligands, such as elongation factors EF-Tu and ET-G (thus inhibiting translation), and endogenous full-length RelA (thus inhibiting (p)ppGpp accumulation upon amino acid starvation, resulting in a relaxed phenotype (Figure 1 B and C)). This interpretation is further substantiated by an earlier report demonstrating that RelA^CTD^ is ribosome-bound (Yang and Ishiguro, 2001). Conversely, at low expression levels, that are more similar to that of the endogenous full-length RelA, expression of neither RelA^CTD^ nor catalytically-inactive G251E mutant RelA* compromises the response of wt MG1655 *E. coli* cells to acute amino acid starvation (Figure 2C and Supplementary Figure 4). We interpret this more physiologically-relevant experiment to mean that neither of the constructs can drive the formation of catalytically-inactive hetero-dimers with native endogenous full-length RelA. Finally, our biochemical experiments using untagged full-length RelA fail to detect reduction in the ppGpp synthetic activity upon increase in enzyme’s concentration that is expected to drive the formation of inactive dimers (Figure 6). Our biochemical results are in good agreement with gel-filtration analysis of full-length *Thermus thermophilus* Rel that failed to detect protein dimerization (Van Nerom et al., 2019). Importantly, the said gel-filtration experiments was performed at unphysiologically high ionic strength that is necessary to keep relatively (in comparison to *E. coli* RelA) soluble *Thermus thermophilus* Rel in solution, and, therefore, should not be over-interpreted. Finally, *Mycobacterium tuberculosis* Rel ectopically expressed in *E. coli* does not form oligomers, further supporting the monomeric nature of RelA/Rel enzymes (Jain et al., 2006).

In addition to characterising the intact RelA regulatory CTD region, we have examined a C-terminally truncated RelA variant lacking the RRM domain (RelA^ΔRRM^). Surprisingly, in the cell, the (p)ppGpp synthetic activity of the enzyme is unregulated in the absence of the RRM domain (Figure 4). Importantly, ectopic expression of RelA^ΔRRM^, is more inhibitory to growth than that of well-characterised constitutively active RelA^NTD^ (Schreiber et al., 1991;Svitil et al., 1993;Gropp et al., 2001). A possible explanation is that while RelA^NTD^ is highly unstable and decays with a half-life of only a few minutes (Schreiber et al., 1991), the stability of RelA^ΔRRM^ is similar to full-length RelA, i.e. half-life of more than two hours (Schreiber et al., 1991). Therefore, more efficient inhibition of growth by RelA^ΔRRM^ is likely is a compound effect of, first, de-regulation of the enzyme leading to uncontrolled (p)ppGpp production similarly to RelA^NTD^, and second, relatively high stability of the protein product. Curiously, while *Francisella tularensis* RelA naturally lacks the RRM domain (Atkinson et al., 2011), the enzyme displays normal, wt-like enzymatic functionality in biochemical assays (Wilkinson et al., 2015). Further work is required to determine the exact regulatory role of the RRM domain that is near-universally conserved in ‘long’ RSHs Rel, RelA and SpoT (Atkinson et al., 2011).

## 6 Conflict of Interest

*The authors declare that the research was conducted in the absence of any commercial or financial relationships that could be construed as a potential conflict of interest*.

## 7 Author Contributions

MR conceived the study, MR and VH designed experiments, KJT, ID, SL and MR performed experiments, contributed to the data analysis, and drafted and revised the manuscript with contributions from VH. All authors have read and approved the final version of this manuscript.

## 8 Funding

This work was carried out at both the Centre for Bacterial Stress Response and Persistence (BASP) at the University of Copenhagen, and the Department of Molecular Biology at Umeå University. The work was supported by a grant from the Danish National Research Foundation (DNFR120), the MIMS Excellence by Choice Postdoctoral Fellowship Programme to M.R. and the Swedish Research council (grant 2017-03783 to VH).

## 9 Acknowledgments

We would like to thank Professor Kenn Gerdes, Dr Anurag Sinha and Dr Farshid Jalalvand for helpful discussions and comments on the manuscript, Dr Szabolcs Semsey for kindly providing plasmid pSEM3034UR2, and the Protein Expertise Platform (PEP) at Umeå University and Mikael Lindberg for constructing pET24d::*His_10_-SUMO* and purifying His_6_-Ulp1. The manuscript was deposited as a preprint to bioRxiv (Turnbull et al., 2019).

## Notes

#### Summary of Updates

Biochemical data added, Figures 5 and 6. The text was revised accordingly throughout.

