## Supplemental Table 1 for "Intramolecular interactions dominate the autoregulation of *Escherichia coli* stringent factor RelA"

### *Supplementary Material*

#### 1 Supplementary Materials and Methods

##### 1.1 Strain construction

MG1655Δ*relA* (Δ*relA*) – the *relA* gene was deleted from the chromosome of MG1655 by P1 transduction as previously described (Sambrook, 1989). Insertion of the kanamycin resistance gene from the Keio collection strain JW2755 (Baba et al., 2006) into correct positions was verified by colony PCR. To remove the Kanamycin resistance cassette from MG1655Δ*relA::kan*, the strain was transformed with the temperature sensitive plasmid pCP20 carrying the yeast FLP recombinase gene (Cherepanov and Wackernagel, 1995). Cells were grown overnight at 30°C with antibiotic selection for the plasmid. Transformants were then re-streaked onto LB agar plates lacking antibiotics and incubated at 42°C overnight. Colonies were then tested for loss of both the Kanamycin antibiotic cassette (LBA supplemented with 25 μg/ml kanamycin, 37°C) and for loss of the pCP20 plasmid (LBA supplemented with Amp, 30°C). Loss of the kanamycin cassette was then verified by PCR. A full list of strains used in this study can be found in Table 1 in the main text.

##### 1.2 Plasmid construction

The coding regions of full length RelA and the C-terminal domain of RelA (codons 405 to 744), along with an optimized Shine-Dalgarno derived from the parM locus of pNDM220 R1 plasmid (Gotfredsen and Gerdes, 1998), were amplified using genomic *E. coli* MG1655 genomic DNA as template and primer sets 1 and 2 or 2 and 3, respectively (Supplementary Table 1). The PCR products were then digested with FastDigest *Eco*RI and *Bam*HI restriction enzymes (ThermoScientific) and cloned into cut pMG25, resulting in vectors pMG25::*relA* and pMG25::*relA*$^{CTD}$, respectively.

pNDM220::*relA*, pNDM220::*relA*$^{\Delta RRM}$, pNDM220::*relA*$^{\Delta ZFD\text{-}RRM}$, pNDM220::*RelA*$^{NTD}$, and pNDM220::*RelA*$^{CTD}$, were constructed by amplifying the corresponding fragments; full-length *relA* (primers 4 and 5), *relA*$^{\Delta RRM}$ (primers 4 and 7), *relA*$^{\Delta ZFD\text{-}RRM}$ (primers 4 and 17), *RelA*$^{NTD}$ (primers 4 and 8), and *relA*$^{CTD}$ (primers 5 and 6), using genomic *E. coli* MG1655 DNA as template. The PCR products were digested with digest enzymes *Bam*HI and *Xho*I (ThermoScientific) and cloned into cut pNDM220.

RelA mutations G251E (primers 9 and 10), C612G (primers 11 and 12), D637R (primers 13 and 14), and C638F (primers 15 and 16) mutation were introduced into the relevant plasmids using two step PCR. A full list of constructed plasmids and their genotypes can be found in Table 1. Oligonuclotides used can be found in Supplementary Table 1.

The wild-type *relA* gene from *E. coli* was amplified using the primer set 17 and 18 (Supplementary Table 1), cloned with *Eco31*I and *Hind*III into pET24d expression vector with N-terminal $His_{10}$-SUMO tag. The expression vector was constructed by the Protein Expertise Platform facility at Umeå University.

### 2 Supplementary Figures and Tables

#### 2.1 Supplementary Figures

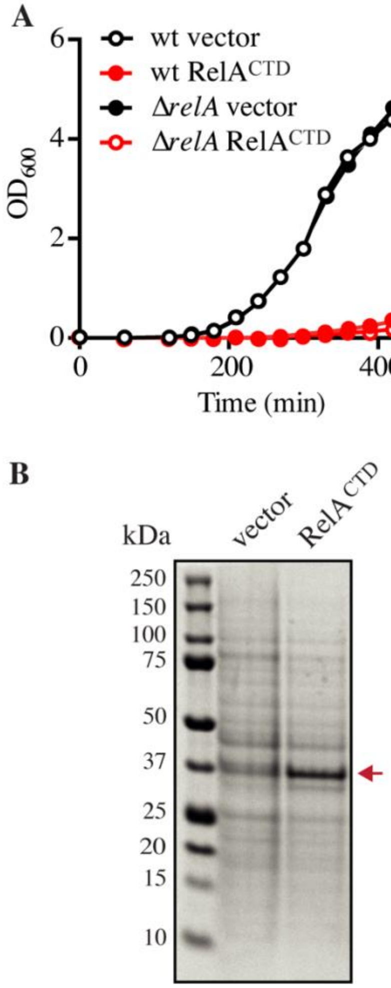


**Supplementary Figure 1. High-level ectopic expression of RelA$^{CTD}$ is inhibitory to growth, related to Figure 1. (A)** Overnight cultures of *E. coli* MG1655 (wt) and MG1655Δ*relA* (Δ*relA*) transformed with pMG25 (vector) or pMG25::*relA$^{CTD}$* (RelA$^{CTD}$), grown in LB with 100 µg/ml ampicillin, were diluted 1/10,000 into fresh LB supplemented with 100 µg/ml ampicillin and 1 mM IPTG, growth was then monitored by sampling at regular time points where $OD_{600}$ measurements were taken. **(B)** SDS-PAGE showing the induction of exponentially growing *E. coli* MG1655 carrying high copy IPTG inducible vector pMG25 (vector) or pMG25::*relA$^{CTD}$* (RelA$^{CTD}$), with 1 mM IPTG for 1 h. RelA$^{CTD}$ is indicated by the red arrow.

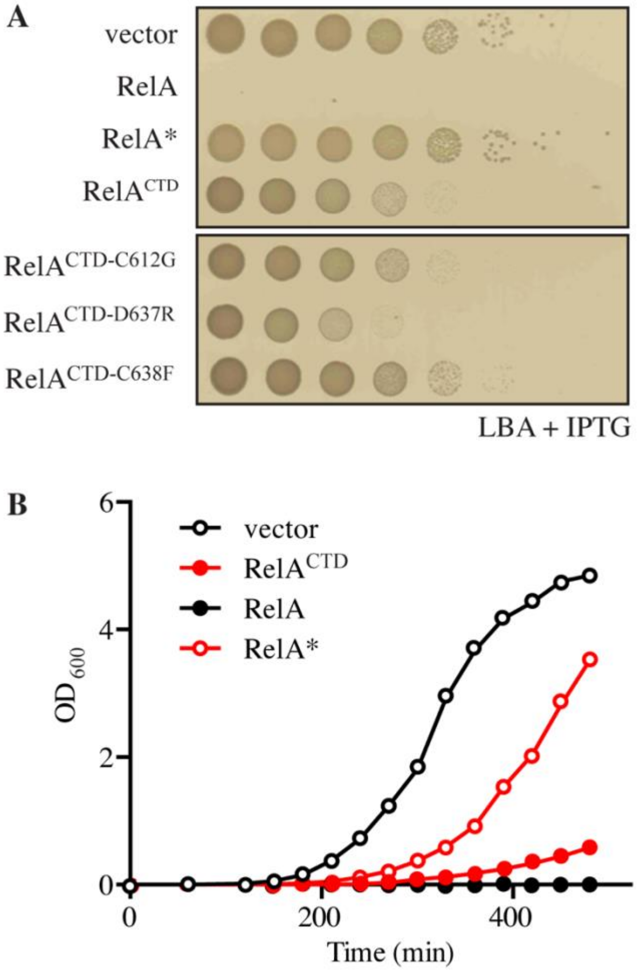


**Supplementary Figure 2. High ectopic expression of RelA, RelA* and RelA$^{CTD}$ inhibit growth, related to Figures 1 and 3. (A)** *E. coli* MG1655 cells were transformed with high copy IPTG inducible vector pMG25 (vector), pMG25::*relA*, pMG25::*relA**, pMG25::*relA*$^{CTD}$, pMG25::*relA*$^{CTD\text{-}C612G}$, pMG25::*relA*$^{CTD\text{-}D637R}$, and pMG25::*relA*$^{CTD\text{-}C638F}$. Ten-fold serial dilutions of overnight cultures grown in LB were prepared and spotted onto LB agar (LBA) supplemented with 100 µg/ml ampicillin and 1 mM IPTG. **(B)** Overnight cultures of *E. coli* MG1655 transformed with pMG25 (vector), pMG25::*relA*, pMG25::*relA**, pMG25::*relA*$^{CTD}$ grown in LB with 100 µg/ml ampicillin, were diluted 1/10,000 into fresh LB supplemented with 100 µg/ml ampicillin and 1 mM IPTG and growth was monitored at $OD_{600}$.

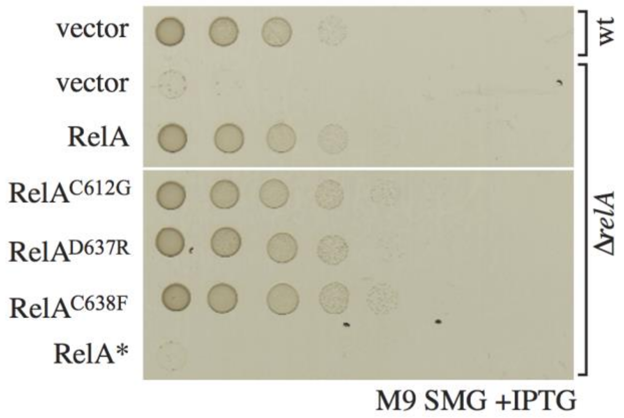


**Supplementary Figure 3. Complementation analysis of wild-type and mutant RelA.** *E. coli* MG1655 cells were transformed with the low copy IPTG inducible vector pNDM220 as positive control (vector, top row). *E. coli* MG1655Δ*relA* (Δ*relA*) cells were transformed with pNDM220 (vector), pNDM220::*relA*, pNDM220::*relA*$^{C612G}$, pNDM220::*relA*$^{D637R}$, pNDM220::*relA*$^{C638F}$, and pNDM220::*relA**. The transformed cells were grown at 37°C overnight in M9 minimal medium with 30 µg/ml ampicillin. Ten-fold serial dilutions were made and spotted onto SMG plate plus 30 µg/ml ampicillin and 1 mM IPTG.

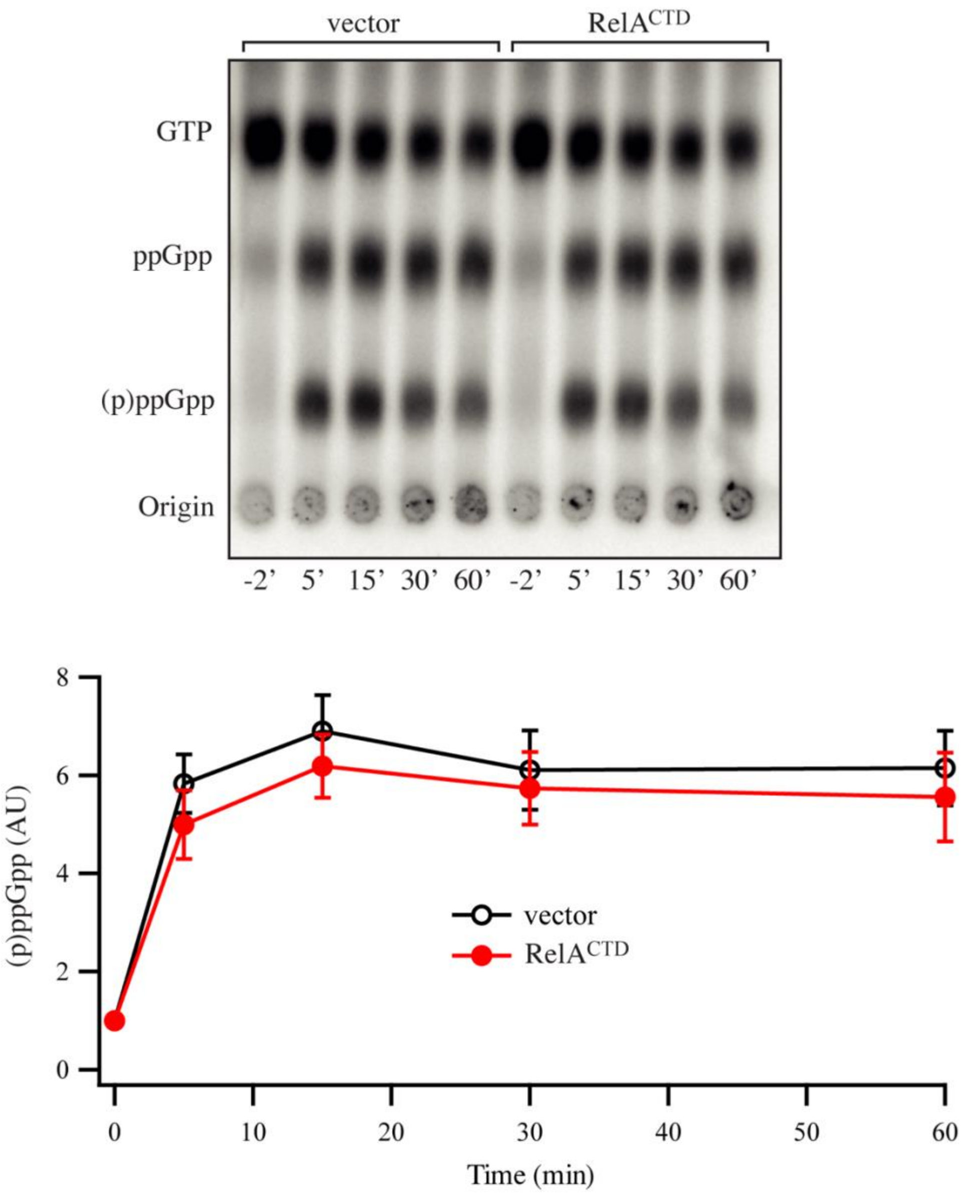


**Supplementary Figure 4. Low-level ectopic expression of $RelA^{CTD}$ does not affect RelA-mediated (p)ppGpp synthesis, related to Figure 2.** Representative audioradiogram of a PEI Cellulose TLC plate showing (p)ppGpp accumulation of *E. coli* MG1655 carrying low copy IPTG inducible vector pNDM220 (vector) or pNDM220::*$relA^{CTD}$* upon serine hydroxamate-induced amino acid starvation. The curves represent the average fold increase for three independent measurements, and the error bars represent standard errors. The levels of (p)ppGpp were normalized to the pre-starved level for each strain. See experimental procedures for more details.

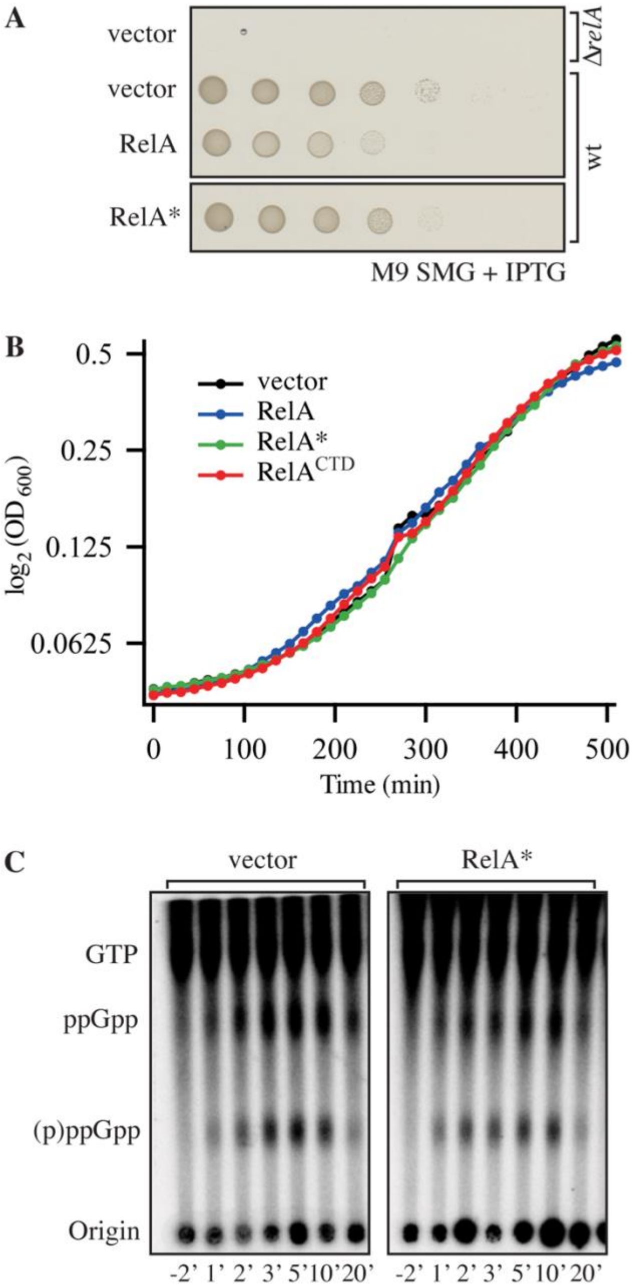


**Supplementary Figure 5. Low-level ectopic expression of RelA* or RelA$^{CTD}$ does not affect RelA-mediated (p)ppGpp synthesis. (A)** *E. coli* MG1655 cells were transformed with low copy IPTG inducible vector, pNDM220 (vector), pNDM220::*relA*, and pNDM220::*relA**. Ten-fold serial dilutions of overnight cultures grown in M9 media, were made and spotted on SMG plate supplemented with 100 µg/ml ampicillin and 1 mM IPTG. MG1655Δ*relA* (Δ*relA*) cells transformed with pNDM220 were used as negative control. **(B)** Overnights of *E. coli* MG1655 cells (grown as described in A), transformed with pNDM220 (vector), pNDM220::*relA*, pNDM220::*relA**, and pNDM220::*relA*$^{CTD}$ were diluted 100-fold and growth was examined in MOPS minimal medium supplemented with SMG (100 µg/ml final), 30 µg/ml ampicillin and 1 mM IPTG. **(C)** Representative audioradiogram of a PEI Cellulose TLC plate showing (p)ppGpp accumulation of *E. coli* MG1655 carrying pNDM220 (vector) or pNDM220::*relA** upon valine-induced isoleucine starvation. See experimental procedures for more details.

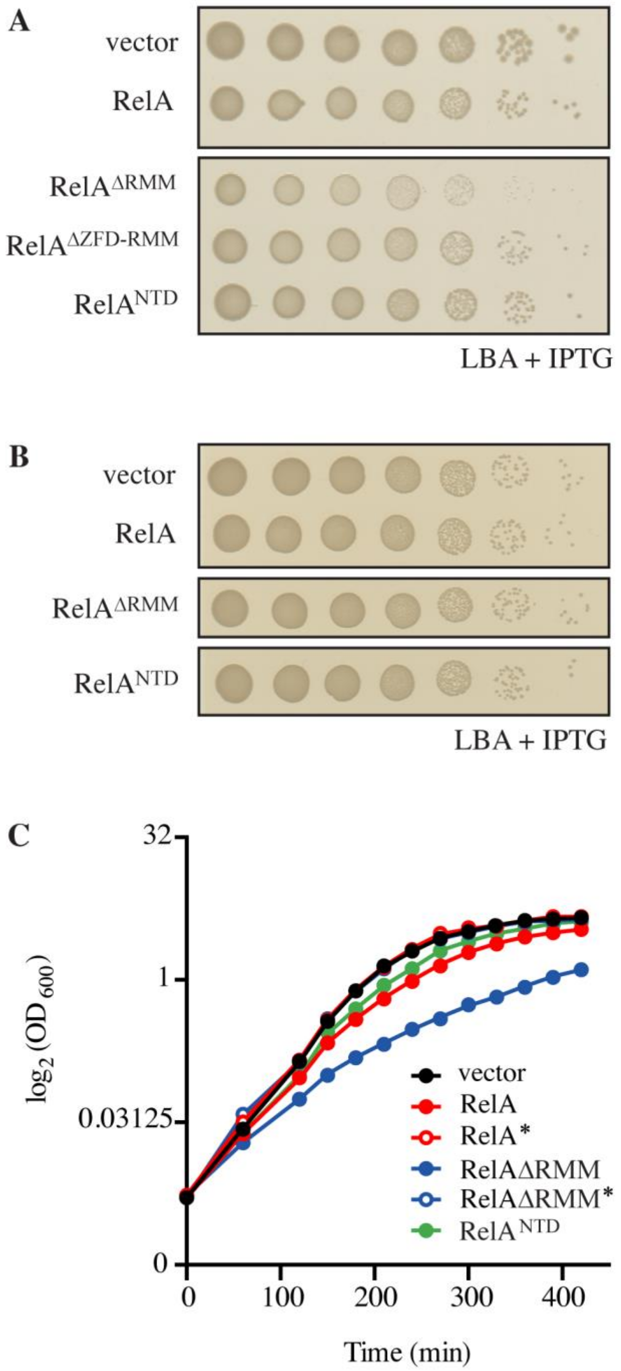


**Supplementary Figure 6. Effects on growth of low-level ectopic expression of RelA variants, related to Figure 4. (A)** *E. coli* MG1655 cells were transformed with low copy IPTG inducible vector, pNDM220 (vector), pNDM220::*relA*, pNDM220::*relA*$^{\Delta RRM}$, pNDM220::*relA*$^{\Delta ZFD\text{-}RRM}$, or pNDM220::*relA*$^{NTD}$. Ten-fold serial dilutions of overnight LB cultures were made and spotted onto LB agar (LBA) supplemented with 30 μg/ml ampicillin and 1 mM IPTG. **(B)** Loading control for Figure 4A in the main text. **(C)** Overnight cultures of *E. coli* MG1655 transformed with low copy IPTG inducible vector, pNDM220 (vector), pNDM220::*relA*, pNDM220::*relA**, pNDM220::*relA*$^{\Delta RRM}$, pNDM220::*relA*$^{\Delta RRM}$*, or pNDM220::*relA*$^{NTD}$, grown were diluted 1/1,000 into fresh LB supplemented with 30 μg/ml ampicillin and 1 mM IPTG and growth was monitored at OD$_{600}$. Doubling times are presented in Figure 4A.

#### 2.2 Supplementary Tables

**Supplementary Table 1.** Oligonucleotides used in this study

| Primer | Sequence |
|---|---|
| 1 | CCCCCGAATTCTAAGGAGTTTTATAAATGGTTGCGGTAAGAAGTGCAC |
| 2 | CCCCCGGATCCTTAACTCCCGTGCAACCGACGCG |
| 3 | CCCCCGAATTCTAAGGAGTTTTATAAATGTACGTCTTTACGCCGAAAGG |
| 4 | CCCCCGGATCCTGGTCCCTAAAGGAGAGG |
| 5 | CCCCCCTCGAGTTAACTCCCGTGCAACCGACGCG |
| 6 | CCCCCGGATCCTGGTCCCTAAAGGAGAGGACGATGTACGTCTTTACGCCGAAAGG |
| 7 | CCCCCCTCGAGTTACCATACCGCGTCAACAATGC |
| 8 | CCCCCCTCGAGTTACAGCTGGTAGGTGAACGGCAC |
| 9 | GCGGAAGTGTATGAACGTCCGAAACACATCTACAG |
| 10 | CTGTAGATGTGTTTCGGACGTTCATACACTTCCGC |
| 11 | CCACATCGCGCGCGGCTGCCAGCCGATTCC |
| 12 | GGAATCGGCTGGCAGCCGCGCGCGATGTGG |
| 13 | GTACACCGCGCCCGTTGCGAACAACTGGCGG |
| 14 | CCGCCAGTTGTTCGCAACGGGCGCGGTGTAC |
| 15 | GTACACCGCGCCGATTTCGAACAACTGGCGG |
| 16 | CCGCCAGTTGTTCGAAATCGGCGCGGTGTAC |
| 17 | CCCCCCTCGAGTTAGCGACCGTTATCTTTACTGCG |
| 18 | GTAACGGTCTCAGGTGGTGTTGCGGTAAGAAGTGCACATATC |
| 19 | GTTACAAGCTTCAACTCCCGTGCAACCG |
